# SpiE is a distinct gut microbial 4-cholesten-3-one 5β-reductase that acts with SpiR to convert cholesterol to coprostanol

**DOI:** 10.64898/2026.10.06.757028

**Authors:** Angela K. Jiang, Gabriela Arp, Margaret Kato, Tess Brunner, Maggie R. Grant, Alexandra M. Clarke, Ashley M. Sidebottom, Mary McMillin, Christopher J. Lehmann, Matthew A. Odenwald, Krysta S. Wolfe, Bhakti K. Patel, Olatoyosi Odenike, Yue Li, Samuel H. Light, Brantley Hall, Xiaofang Jiang

**Author notes:** Corresponding authors: Brantley Hall and Xiaofang Jiang. These authors contributed equally (Angela K. Jiang, Gabriela Arp).

## Abstract

Gut microbes convert cholesterol into coprostanol, a poorly absorbed sterol that is largely excreted in feces and is linked to lower serum cholesterol levels. However, the enzymes responsible for this process remain poorly characterized. The second step of the pathway, cholestenone reduction into coprostanone, has long been established, yet the enzyme(s) responsible were unknown. Here, we use comparative genomics to identify SpiE, a cholestenone (4-cholesten-3-one) reductase that is part of the steroid processing (*spi*) pathway with SpiR. We confirm that SpiE binds to and reduces both cholestenone and progesterone into 5β-reduced forms. SpiR and SpiE together convert cholesterol into coprostanol, indicating that both enzymes are sufficient for coprostanol formation. SpiE belongs to the short-chain dehydrogenase (SDR) family and represents an evolutionarily distinct lineage of SDR ene-reductases. Metagenomic analyses reveal that *spiE* relative abundance is strongly correlated with stool coprostanol (Spearman’s ρ=0.81, *p*=5.99e-65). In longitudinal patient cohorts, antibiotic treatment coincided with loss of SpiE and coprostanol production. Post antibiotics, the emergence of distinct SpiE-encoding genera rescued coprostanol production in some individuals. Our findings provide the biochemical basis of the cholesterol to coprostanol pathway and lay the groundwork for future studies linking it to host physiology.

## Introduction

The human intestinal lumen is a densely populated ecosystem, in which microbes compete to ferment undigested, soluble fibers while growing in the absence of molecular oxygen^1^. Gut microbes rely on alternative electron acceptors and fermentative pathways to maintain intracellular redox balance^2^. High-energy electrons carried by reduced cofactors generated during fermentation are essential for downstream reductive processes, while substrate-level phosphorylation provides ATP, making reductive and oxidative enzymatic reactions central processes within the gut^3^.

Although microbial steroid transformations have traditionally been studied for their effects on host physiology^4^, host-derived steroids may be among the most abundant metabolites used by the gut microbiota as terminal electron acceptors that support energy conservation and growth. In the anoxic colonic environment, microbial oxidoreductases can couple the reduction of steroids with the oxidation of cofactors like NADH, regenerating NAD+ and maintaining cellular redox balance^5–8^. This suggests that steroid metabolism serves as a component of anaerobic energy metabolism. In support of this idea, *Eubacterium sp.* ATCC 21408, a gut bacterium capable of converting cholesterol to coprostanol, requires cholesterol for growth, suggesting steroid metabolism is involved in cellular ATP generation^9^.

Among these steroids is cholesterol, a central steroid hormone precursor, which enters the intestinal lumen at 1 gram per day from a combination of diet, bile, and epithelial turnover; 50-60% is reabsorbed, while the remaining is available for microbial metabolism^4,10^. Gut microbes convert cholesterol into coprostanol, a poorly absorbed sterol that can be readily excreted^11^. Fecal coprostanol-to-cholesterol ratios correlate inversely with serum cholesterol, implying that gut microbial conversion of cholesterol is associated with host cholesterol homeostasis^12^. The microbial basis of this process is supported by multiple lines of evidence. Germ-free animals fail to produce coprostanol and display elevated serum cholesterol levels^13,14^, while antibiotic treatment similarly impairs conversion and raises cholesterol levels^15^, and increased abundance of gut coprostanoligenic bacteria correlates with both greater conversion efficiency and lower circulating cholesterol^16,17^. These observations suggest that microbial cholesterol metabolism may contribute to cholesterol homeostasis, and highlight the need to understand its underlying microbial enzymatic mechanisms.

Cholesterol metabolism by gut bacteria proceeds through three sequential reactions. First, cholesterol is oxidized and Δ^5–4^ isomerized into cholestenone. In the second step, an unknown 5β-reductase reduces cholestenone (4-cholesten-3-one) into coprostanone. Finally, coprostanone is reduced into coprostanol (**Figure 1b**)^18^. The enzymes responsible for the conversion of cholesterol into coprostanol have been partially characterized. The IsmA enzyme in *Eubacterium coprostanoligenes* ATCC 51222 mediates cholesterol oxidation and coprostanone reduction^17^. Our group subsequently identified SpiR (<u>S</u>terol <u>p</u>rocessing <u>i</u>somerase and <u>R</u>eductase), a stronger predictor of gut microbial cholesterol conversion, as a gut microbial enzyme that catalyzes the first step, the oxidation and Δ^5–4^ isomerization of cholesterol into cholestenone^6^, while reducing NAD+ to NADH. This enzyme is part of the short chain dehydrogenase family and is lineage-specific to a clade of uncultured gut bacteria in the family Acutalibacteraceae. Combined with the final two reduction steps into coprostanol, cholesterol conversion is a net electron sink for the cell.

**Figure 1:**
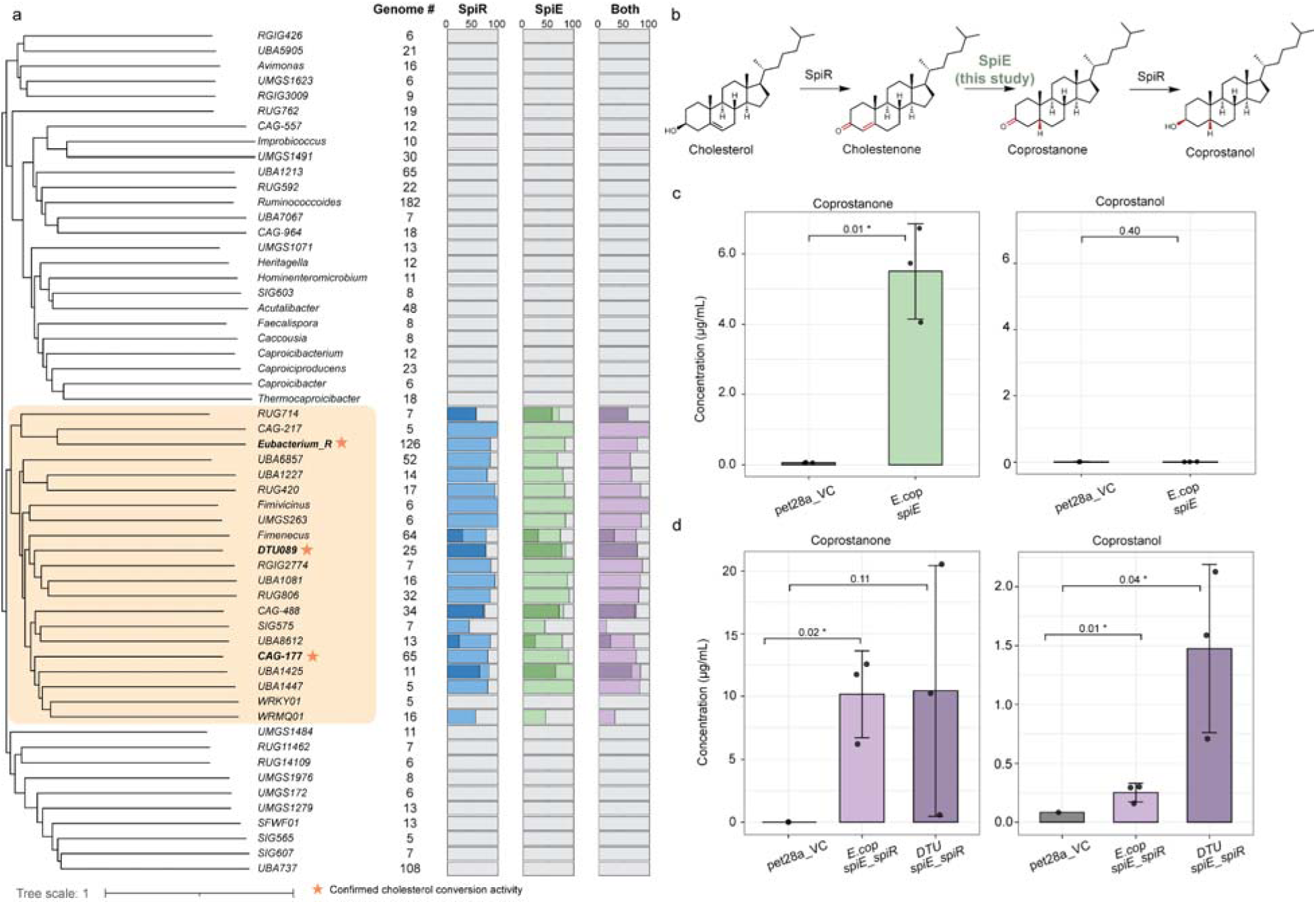
a) Phylogenetic distribution of SpiR and SpiE homologs across Acutalibacteraceae genera. The phylogram shows the taxonomic relationships among genera in the family Acutalibacteraceae. Genera with fewer than five representative genomes are not displayed. The number of genomes screened per genus and the percentage encoding SpiR, SpiE, and both are shown on the right. Darker bars indicate the genes were found within the same neighborhood (defined as ±5000 base pairs from *spiE*). Genera containing species with confirmed cholesterol conversion activity, either detected directly in the organism or validated through heterologous expression of the enzyme in *E. coli* (this study), are labelled with stars. b) Enzymatic pathway of *E. coprostanoligenes* ATCC 51222 conversion of cholesterol into coprostanol via SpiR and SpiE. SpiE reduces cholestenone into coprostanone. c) Quantification of coprostanone and coprostanol production by *E. coli* expressing SpiE_ecop_ when incubated with cholestenone. d) Quantification of coprostanone and coprostanol production by *E. coli* expressing SpiE_ecop_ and SpiR_ecop_ incubated with cholesterol, and SpiE_DTU089_ and SpiR_DTU089_. For c) and d), each point represents one biological replicate. The bar represents the mean of 3 biological replicates, and the error bars represent standard error. The *p*-values represent the results of one-sided Welch’s t-tests.

The enzyme catalyzing the first of the two reductions in the cholesterol-to-coprostanol pathway, the conversion of cholestenone to coprostanone, has remained unidentified. We now report SpiE as a cholestenone (4-cholesten-3-one) 5β-reductase responsible for this step. Through heterologous expression and GC-MS, we confirm that SpiE reduces cholestenone into coprostanone and progesterone into 5β-dihydroprogesterone. SpiE coexpressed with SpiR converts cholesterol into coprostanol, implicating SpiR in the reduction of coprostanone into coprostanol. Phylogenetically, SpiE represents a distinct lineage of ene-reductases in the short chain dehydrogenase family and contains a distinct active site architecture. Finally, metagenomic analyses reveal that *spiE* is correlated with coprostanol detection in stool samples across four cohorts, establishing it as a biomarker for cholesterol conversion. Within antibiotic-treated individuals, SpiE depletion and recovery coincided with coprostanol production loss and rescue. Together with SpiR, the discovery of SpiE accounts for the three enzymatic steps of gut microbial cholesterol-to-coprostanol conversion, a pathway first observed in the 1930s^19–21^.

## Results

### SpiE is a putative cholestenone 5β ene-reductase in *Eubacterium coprostanoligenes*

In our previous study, we identified SpiR as an enzyme responsible for catalyzing the reaction of cholesterol into cholestenone, the first step in cholesterol conversion by gut bacteria^6^. Given that the conversion of cholesterol to coprostanol is a 3-step multi-enzyme pathway, we hypothesized that the missing 5β ene-reductase involved in the second step of the conversion of cholesterol into coprostanol may have co-evolved with SpiR. We also further hypothesized that this 5β ene-reductase, like SpiR, is lineage-specific to a clade in Acutalibacteraceae. Therefore, we sought to identify gene families that cooccur with SpiR within Acutalibacteraceae.

We used Evolink to identify OrthoFinder-inferred Acutalibacteraceae gene families coevolved with SpiR^22^. The tool calculates an Evolink index, which is a measure of phylogenetic co-occurrence ranging from -1 to 1, where values near 1 indicate identical genotype and phenotype distributions across the phylogeny, and sets a significance threshold using Isolation Forest detection of scores deviating from the background distribution. We identified 4 gene families significantly positively correlated with SpiR, not including SpiR itself (**Figure S1**). Of those, the gene family most strongly correlated to SpiR was annotated as a putative NAD(P)-dependent oxidoreductase in the short chain dehydrogenase (SDR) family (IPR051783) with a Rossmann-fold domain (Evolink index: 0.78, Score: 0.87). This enzyme is also present in the genomic neighborhoods of 17.5% *spiR* homologs (**Figure 1a**). Unexpectedly, this enzyme is not part of the OYE family, which typically catalyze steroid ene-reductions^7,23,24^. Although SDRs are primarily associated with the reduction of ketones and aldehydes, several plant short chain dehydrogenase family enzymes participate in steroid ene-reduction^25,26^. We therefore hypothesized that this gene family encodes cholestenone reductase, and named it **S**terol **p**rocessing **i**ntestinal **E**ne-reductase (SpiE).

### SpiE catalyzes the conversion of cholestenone into coprostanone

To test the activity of SpiE, we heterologously expressed SpiE of *E. coprostanoligenes* ATCC 51222 (SpiE_ecop_) in media with cholestenone. The identities of the products were confirmed using gas chromatography-mass spectrometry (GC-MS). *E. coli* expressing SpiE_ecop_ showed conversion of cholestenone into coprostanone (mean 5.51 µg/ml) whereas the vector controls did not, indicating cholestenone reduction activity in SpiE (**Figure 1c**). There was no difference between coprostanol production in the vector control and *E. coli* expressing SpiE, indicating that SpiE does not reduce the 3-oxo group on coprostanone (**Figure 1c**). These results show that SpiE confers 5β ene-reductase activity on cholestenone.

To test whether SpiE can reduce a broader range of Δ^4^-3-ketosteroid substrates, we heterologously expressed SpiE_ecop_ in *E. coli* and incubated the cells with progesterone, and then used LC-MS to detect products. Consistent with our cholestenone observations, we detected 5β-dihydroprogesterone in *E. coli* expressing SpiE_ecop_ and not in the vector controls (**Figure S4b**). Epipregnanolone was not detected, consistent with the cholestenone results (**Figure S4c**). This confirms that SpiE is able to perform 5β-reduction on a variety of steroid substrates.

### SpiR and SpiE are sufficient for coprostanol formation from cholesterol

We have previously shown that SpiR catalyzes the first step in the oxidation and isomerization of cholesterol into cholestenone. To determine whether SpiE and SpiR are sufficient for coprostanol production, we heterologously expressed SpiE_ecop_ and SpiR_ecop_ together in *E. coli* cultured in media containing cholesterol. We also expressed *spiR* and *spiE* homologs identified in an uncultured Acutalibacteraceae species *DTU089 sp015066225* (NCBI accession: GCA_015066225.1). These homologs (SpiR_DTU089_ and SpiE_DTU089_) share 57% and 65% protein identity to SpiR_ecop_ and SpiE_ecop_, respectively, and are adjacent and co-oriented in the genome. We quantified coprostanone and coprostanol using GC-MS.

*E. coli* expressing SpiE_ecop_ and SpiR_ecop_ in cholesterol media produced coprostanone (mean of 10.17 µg/ml across three replicates), and coprostanol (mean of 0.25 µg/ml), indicating that these two enzymes are sufficient to complete cholesterol reduction into coprostanol (**Figure 1d**). Coprostanone and coprostanol were also detected in *E. coli* expressing SpiE_DTU089_ and SpiR_DTU089_ (mean of 10.46 µg/mL coprostanone, and 1.47 µg/mL coprostanol) (**Figure 1d**). SpiE_ecop_ alone with cholestenone produced coprostanone but no coprostanol above the vector control, which rules out an endogenous *E. coli* 3-oxo reductase (**Figure 1c**). Therefore, the coprostanol produced in SpiE and SpiR experimental groups is attributable to SpiR. These results indicate that SpiR and SpiE are sufficient for coprostanol production, and that SpiR is implicated in the third step, the reduction of coprostanone into coprostanol.

These patterns are held in a different steroid substrate. When incubated with pregnenolone, *E. coli* expressing both SpiE_ecop_ and SpiR_ecop_ showed conversion to epipregnanolone (**Figure S4c**). This implicates the *spi* pathway in broader steroid metabolism in the gut.

### SpiE belongs to a distinct class of ene-reductases

SpiE represents an unusual case of a SDR performing ene-reduction. While SDRs are well-characterized for dehydrogenase reactions, ene-reductases that act on alkene steroid substrates are uncommon and poorly characterized in bacteria. To place SpiE in the context of SDR evolution, we constructed a phylogeny of a curated set of SwissProt SSF51735 superfamily enzymes classified as oxidoreductases (EC 1.-.-.-) (**Figure S2, Supplementary Table 2**). Notably, SpiE showed no close phylogenetic relationships to other SDR ene-reductases (EC 1.3.-.-), and instead clustered with enzymes acting on CH-OH groups (EC 1.1.-.-) (**Figure S2**). In addition, SpiE was unrelated to the plant *Digitalis lanata* progesterone 5β-reductase by sequence identity (UniProt Accession: Q6PQJ9) (18.00% amino acid identity, TM-score: 0.64) (**Figure S2**), suggesting convergent evolution of the 3-oxo-Δ^4,5^-steroid 5β-reductase function. The closest SwissProt homolog was Rv1106c, a cholesterol dehydrogenase from *Mycobacterium tuberculosis* (27.3% identity). The closest PDB structural homolog was the SDR family protein Gox2253 (e-value 6.48e-18), an aldehyde reductase. These analyses suggest that SpiE represents a distinct lineage of steroid ene-reductases in bacteria.

To investigate whether SpiE shows a similar mechanism to other SDR enzymes, we identified the catalytic residues (see Methods). Four residues in the putative active site, D77, R79, S118, and Y138, showed total conservation across 485 SpiE homologs (**Figure 2a**). These residues differ from the canonical catalytic tetrad found in other SDR enzymes (Asn-Ser-Tyr-Lys)^27^, possibly underlying SpiE’s ene-reductase activity, which is distinct from the dehydrogenase activities more commonly characterized in this family. To assess the role of these residues in catalysis, we generated alanine substitutions at each position and tested reductase activity using progesterone, which we previously confirmed as a substrate for SpiE (**Figure S4b**). R79A and D77A mutants showed nearly no reductase activity, implicating them as catalytic residues, whereas the S118A and Y138A mutants showed reduced but detectable activity compared to the wild-type, implicating their role in binding (**Figure 2b**).

**Figure 2:**
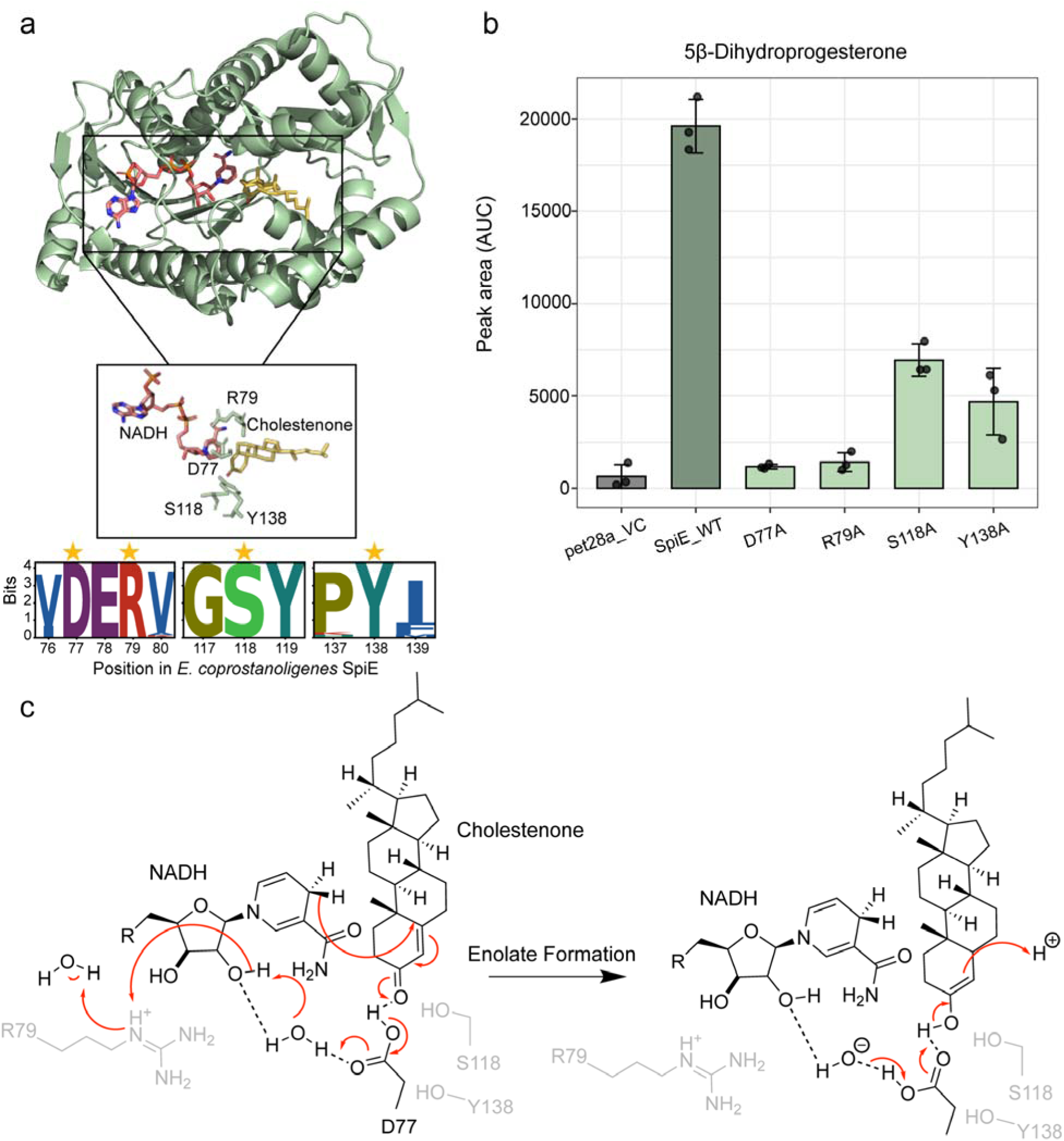
a) Cholestenone and NADH co-folded to the alphafold3-predicted SpiE structure. The predicted interactions of the catalytic residues with cholestenone are shown. b) Bar graph shows the normalized area under the curve of 5β-dihydroprogesterone detected in the identified enzymes across different mutations, indicating the product abundance. Data are presented as mean values ± standard error. c) Proposed enzymatic mechanism of SpiE reduction of cholestenone via enolate formation based on the isopiperitenone reductase mechanism^29^.

We determined SpiE’s preferred cofactor using a biological activity assay with purified SpiE comparing its use of NADH to NADPH **(Figure 3b)**. By the final measurement, the absorbance of SpiE samples incubated with NADH and cholestenone had declined to about 50% of initial value, compared with 80% remaining for NADPH and cholestenone, indicating that SpiE preferentially utilizes NADH over NADPH as a reducing cofactor for cholestenone reduction.

**Figure 3:**
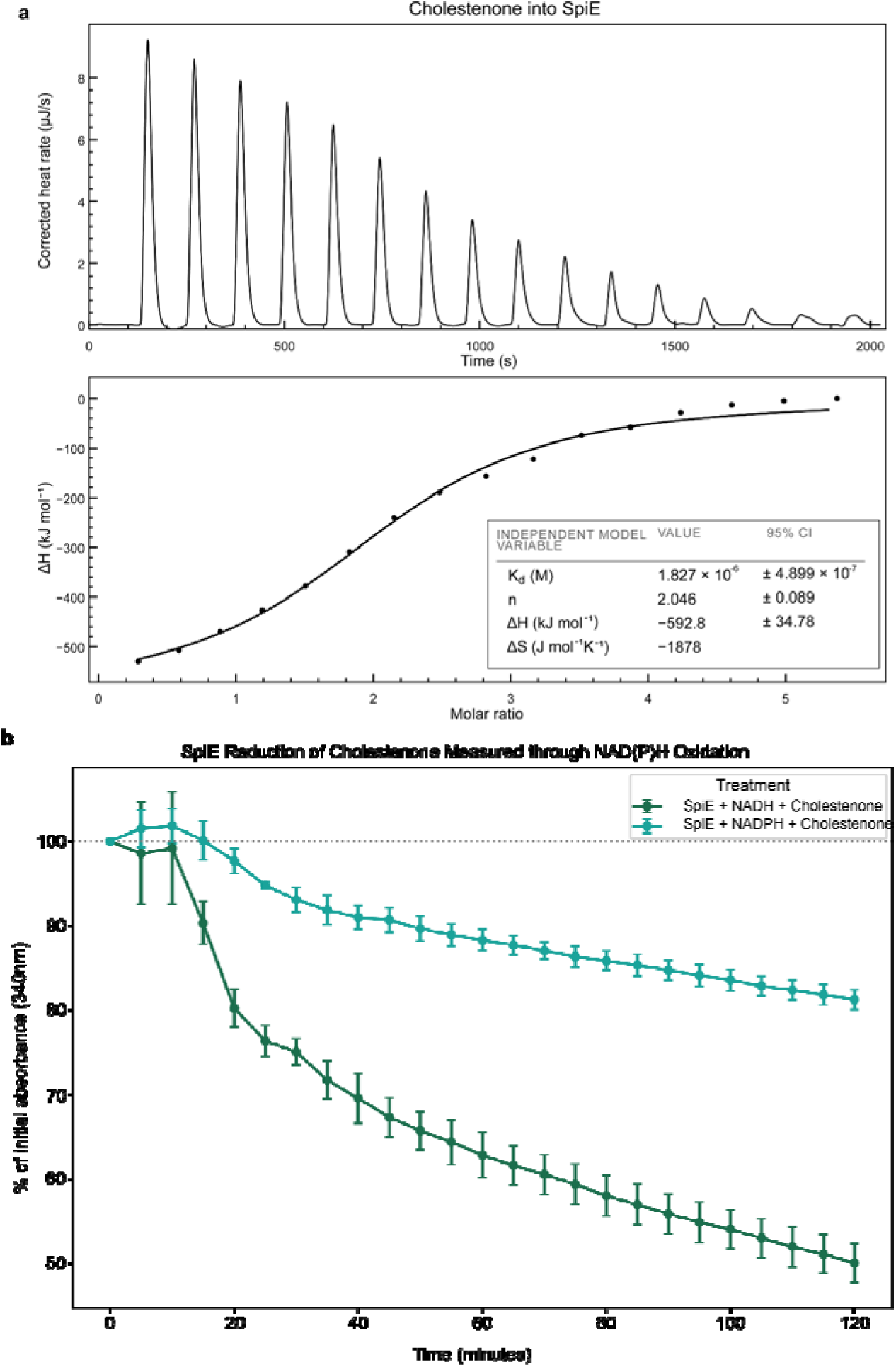
a) Binding of SpiE to cholestenone as measured by ITC. Raw data is shown in the upper graph of the panel, integrated binding isotherms with their independent binding model fitting are shown in the lower panel. The binding of the trace was performed using a 100 µM cholestenone in the syringe and 10 µM SpiE in the calorimeter cell. b) Cholestenone reduction was indirectly monitored by NAD(P)H oxidation at 340 nm, buffer-corrected OD_340_. SpiE + NADH + cholestenone (green) and SpiE + NADPH + cholestenone (turquoise) were compared for cofactor preference. To control for differences in starting NAD(P)H concentration across biological replicates, each replicate was normalized to its own time-zero (T0) absorbance and expressed as a percentage of initial value. Data points represent the mean ± SEM of normalized values from n = 3 independent replicates per condition.

Based on our mutagenesis studies, computational structural predictions of SpiE, and previous studies on SDR ene-reductases, we propose that SpiE follows a similar enolate formation mechanism as observed in other ene-reductases in the SDR family (**Figure 2c**). A previous study showed that the mechanism of cholestenone reduction by a partially purified 5β-reductase involved the transfer of the 4B hydrogen of NADH to the 5β position of cholestenone^28^. D77 is possibly key for catalysis, initiating a proton transfer cascade by obtaining a proton from a nearby water molecule^29^. In isopiperitenone reductase (IPR), this role is performed by glutamic acid. R79 may stabilize the cofactor and promote proton transfer, a role performed by lysine in IPR. Active site residues S118 and Y138 are possibly responsible for stabilizing the enolate intermediate. In IPR, this role is performed by a serine residue.

### SpiE is a homodimer that binds to cholestenone

To investigate the oligomeric state of heterologous SpiE, a dynamic light scatterer (DLS) analysis was performed. The measured hydrodynamic radius was 3.2 nm, which is consistent with the theoretical radius of its dimeric state (3.12 nm) and larger than the expected radius of its monomeric form (2.75 nm). This indicates that SpiE is predominantly present as a dimer in the solution (**Figure S5**). Based on isothermal titration calorimetry (ITC), cholestenone and progesterone exhibited stoichiometric values (n) of 2.046 ± 0.089, 2.423± 0.058, suggesting two ligands bound per SpiE dimer and indicating one binding site per SpiE (**Figure 3a; Figure S3**). The stoichiometry and DLS results suggest that SpiE functions as a homodimer.

We investigated the substrate specificity of SpiE by evaluating its binding affinity to cholestenone and progesterone. ITC experiments also revealed micromolar binding affinity for cholestenone (Kd 1.827E-6, 95% CI: ± 4.899E-7 M) (**Figure 3a**). Sub-micromolar binding was also observed between SpiE and progesterone (Kd: 1.124E-7, 95% CI: ± 4.865E-8) **(Figure S3)**. The results indicate that SpiE and SpiR bind complementary metabolites needed to complete the gut bacteria cholesterol metabolism pathway.

### SpiE is strongly correlated with cholesterol conversion across human gut microbiome cohorts

Our previous work showed that SpiR is positively correlated with cholesterol conversion in stool samples^6^. Because SpiR and SpiE together complete the cholesterol conversion pathway, we hypothesized that SpiE shows similar correlations to cholesterol conversion. To link SpiE to cholesterol metabolism in the gut, we analyzed paired metabolomics and metagenomics data from four cohorts: acute myeloid leukemia (AML, n = 13), liver disease (LD, n = 153), medical intensive care unit (MICU, n = 52), and liver transplant (LT, n = 56) (**Table S3**). Analyzing a single fecal sample per subject, we measured coprostanol, the final product in the cholesterol conversion pathway, using GC-MS, defining cholesterol converters as individuals with coprostanol normalized area under the curve (AUC) greater than 0.1.

SpiE homolog detection was consistently correlated with cholesterol conversion. Across the four cohorts, 35% (96/274) of individuals were cholesterol converters, consistent with previous studies finding that the prevalence of cholesterol converters was variable^30^. 81% (78/96) of cholesterol converters harbored SpiE homologs, compared to 6% (10/178) of non-converters. The proportion of samples with SpiE homologs (CPM ≥ 1) was significantly higher in cholesterol converters from all cohorts (Fisher’s Exact Test; AML: *p* = 0.019; LD: *p* = 8.5 × 10^-20^; MICU: *p* = 1.98 × 10^-10^; LT: *p* = 3.5 × 10^-5^) (**Figure 4a**), and coprostanol abundance was significantly enriched in SpiE-positive samples for all cohorts using a Mann-Whitney U test (AML: U = 39, *p* = 0.045; LD: U = 4235, *p* = 7.2 × 10^-20^; MICU: U = 655, *p* = 1.3 × 10^-8^; LT: U = 517, *p* = 2.7 × 10^-5^) (**Figure 4b**). In addition, SpiE homolog abundance was positively correlated with coprostanol abundance. The Spearman’s correlation coefficient (ρ) when combining all cohorts was 0.81 and the *p*-value was 5.99e-65, indicating strong positive correlation. When stratified by cohort, ρ ranged from 0.78 for the LT and LD cohorts to 0.95 for the AML cohort, with all *p*-values < 0.001 (**Figure S6**). Finally, to assess the strength of SpiE abundance for predicting fecal coprostanol production, we performed receiver operating characteristic (ROC) curve analysis. The resulting model had an area under the curve (AUC) of 0.92, indicating strong classification performance and robustness to variations in detection thresholds (**Figure S7**). Our results demonstrate that SpiE, like SpiR, is strongly associated with the presence of cholesterol metabolism products in stool samples.

**Figure 4:**
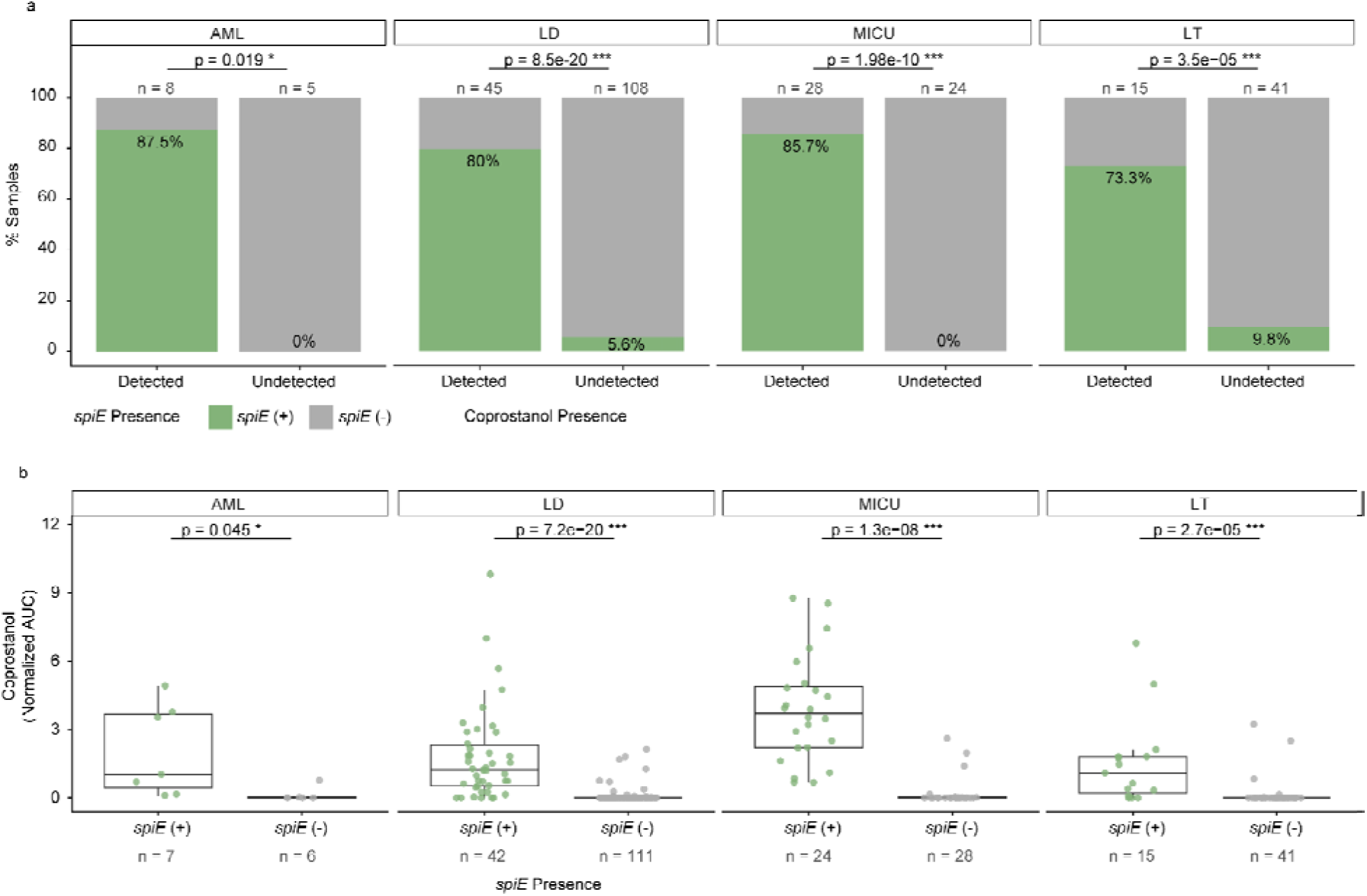
a) Association between SpiE homolog presence and coprostanol presence in four paired human metagenomic and metabolomic datasets. Significant differences in *spiE* detection in coprostanol(+) and coprostanol(-) samples were determined using a Fisher’s exact test with Bonferroni correction. *n* represents the number of coprostanol(+) and coprostanol(-) metagenomic samples analyzed per cohort. b) Association between *spiE* homolog presence and coprostanol abundance in four paired human metagenomic and metabolomic datasets. Coprostanol abundance is measured as normalized area under the curve (AUC). Each point represents a paired metabolomics–metagenomic sample, with one sample per patient. The lines within each box represent the median, the boxes represent the IQR, and the whiskers represent 1.5 × the IQR. Significant differences in coprostanol abundance between *spiE*(+) and *spiE*(-) samples were determined using a Mann-Whitney U test with Bonferroni correction.

### Within-patient loss and replacement of *spiE*-encoding genera tracks coprostanol production over time

Having established the relationship between *spiE* and coprostanol production across patients, we next asked whether the same association holds within individuals over time. We selected thirteen patients that exhibited variable coprostanol levels across the sampling period and performed metagenomic read recruitment to quantify *spiE* abundance at each timepoint. These analyses revealed that *spiE* and coprostanol rose and fell together within serial samples from individual patients, recapitulating the between-individual association at the level of repeated observations from a single host (**Fig. 5**).

**Figure 5:**
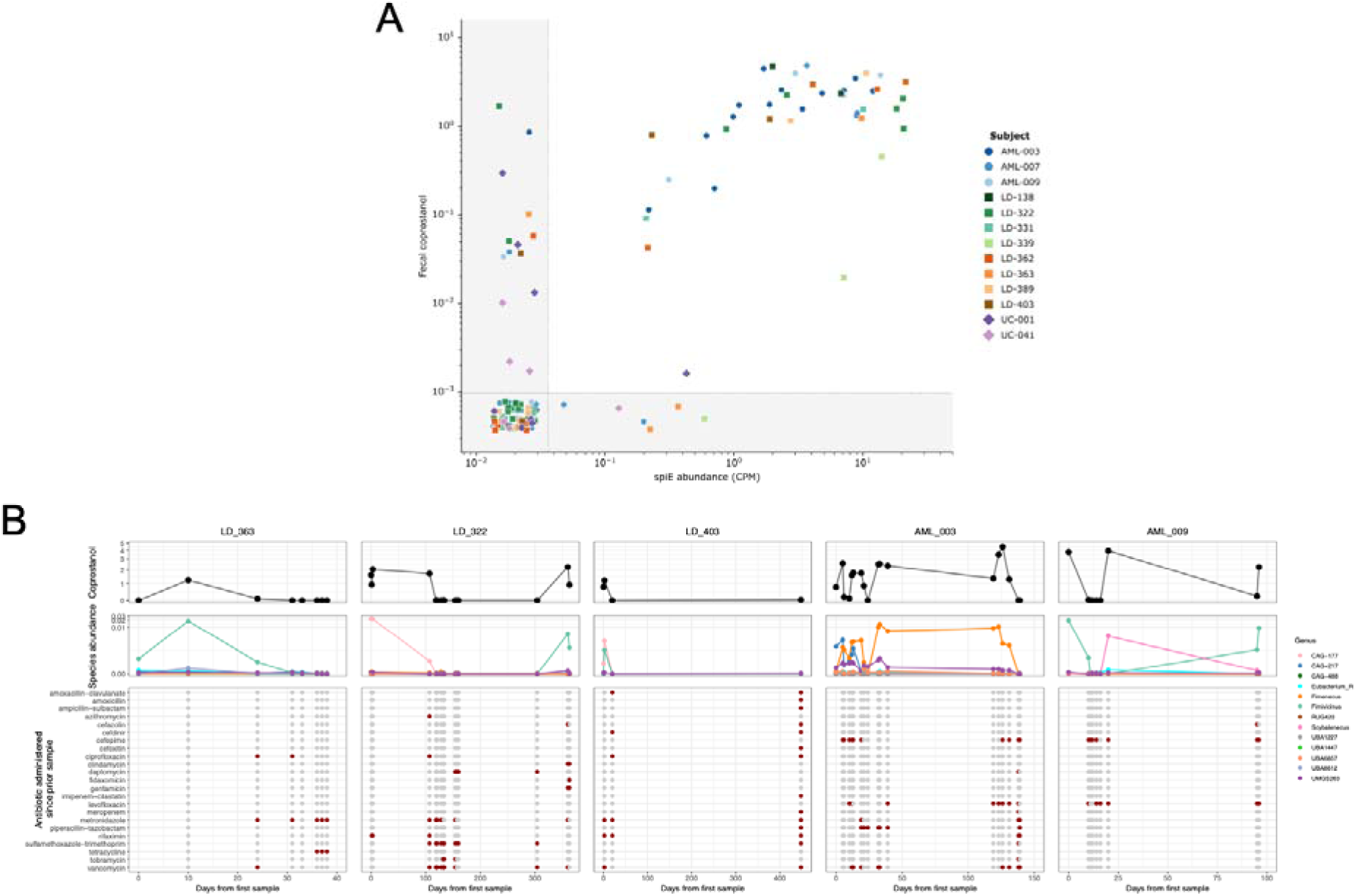
*spiE* abundance predicts fecal coprostanol production within individual patients over time. **(A)** Paired *spiE* abundance and fecal coprostanol across 130 longitudinal specimens from the 13 patients that contributed both *spiE*-positive and *spiE*-negative samples. Shaded margins denote censored observations, comprising samples in which *spiE* read recruitment was zero and samples in which coprostanol fell below the limit of detection (horizontal band). **(B)** Longitudinal profiles for five representative patients, plotted against days elapsed from each subject’s first specimen. Upper track, fecal coprostanol. Middle track, relative abundance of *spiE*-encoding species aggregated by genus. Lower track, antibiotic agents administered in the interval since the preceding specimen, where grey points mark collected specimens and filled points mark administration of the indicated agent.

Further analysis of individual time courses revealed that loss of coprostanol production often coincided with patient antibiotic exposure, consistent with previous reports of the antibiotic susceptibility of coprostanol-producing strains^31^. Notably, following this loss, neither *spiE* nor coprostanol was recovered from any subsequently collected fecal sample in some patients (e.g., AML-003, LD-363, LD-403) (**Fig. 5**). By contrast, coprostanol production resumed at later timepoints in other patients (e.g., LD-322, AML-009) (**Fig. 5**). Notably, this recovery of coprostanol was in some cases associated with metagenomic reads recruiting to *spiE* variants from a bacterial genus distinct from those detected before the loss, suggesting that coprostanol production can be rescued by later acquisition of new coprostanol-producing genera.

## Discussion

In this study, we identify SpiE as a cholestenone 5β-reductase, and implicate SpiR, previously characterized only as a cholesterol dehydrogenase, in the catalysis of the terminal reduction of coprostanone to coprostanol. The strong correlation of SpiE with stool coprostanol levels, both across individuals and within individuals over time, highlights its utility as a metagenomic biomarker for gut bacterial cholesterol conversion. The cholestenone ene-reductase has remained elusive prior to this study. The identification of SpiE, together with the confirmation that SpiE and SpiR together convert cholesterol into coprostanol, resolves the enzymatic basis of the microbial cholesterol to coprostanol pathway, and lays the foundation for future studies on cholesterol metabolism in the gut and how it relates to human health.

Our results support the three-step route from cholesterol to coprostanol: oxidation and Δ^5–4^ isomerization of cholesterol to 4-cholesten-3-one by SpiR, 5β-reduction of 4-cholesten-3-one to coprostanone by SpiE, and reduction of the 3-oxo group of coprostanone to coprostanol by SpiR. A recent preprint proposed an alternative route proceeding through 5-cholesten-3-one with the discovery of IsmB^24^. Our data are instead consistent with earlier isotope-labeling and intermediate-detection studies that widely support 4-cholesten-3-one as the intermediate^18,28,32^. Furthermore, SpiE prefers NADH as a cofactor, matching the requirement reported for a cholestenone 5β-reductase partially purified from rat caecal contents in 1973^28^. Our proposed mechanism involves the transfer of the 4B hydride of NADH to the β-carbon (C5) of the α,β-unsaturated 3-ketone. Like the canonical Old Yellow Enzyme (OYE) mechanism^33^, in which the hydride is instead derived from reduced FMN, this requires an electron-withdrawing group (EWG). The EWG pulls electron density away from the β-carbon, giving it a partially positive charge (δ+), thus promoting nucleophilic attack. 5-cholesten-3-one has an unconjugated Δ5 alkene and is therefore unlikely to be substrate of SpiE and other canonical OYEs. In 2026, the genome of a coprostanoligenic human gut isolate belonging to the Acutalibacteraceae genus *CAG-177* was sequenced^34^. A BLASTp search revealed that its genome contained SpiR_ecop_ (58.6% protein identity) and SpiE_ecop_ (67.4% protein identity) homologs (**Table S1**), but no IsmA homolog and a weak IsmB match (30.5% protein identity). A coprostanol producing isolate therefore contained the entire *spi* pathway but not the *ism* pathway, suggesting that the *ism* pathway is not strictly required for coprostanol production in the human gut.

A variety of steroid 5β-reductases have been characterized, including the flavin-dependent oxidoreductases OsrB^8^, ci2350 in *C. innocuum*^7^, 5β-reductase in *Bacteroides thetaiotaomicron*^35^, and in this present work, SpiE. These reductases share minimal sequence similarity, suggesting convergent evolution. It may be biochemically straightforward to convert other enzyme types into ene-reductases^29^. Gut bacteria have evolved diverse ene-reductases for anaerobic respiration^2^, suggesting that ene-reduction may represent a strategy for accepting reducing equivalents under anaerobic conditions. The repeated presence of steroid ene-reductases may reflect the selection for enzymes that provide steroid substrates for microbial redox metabolism, meaning that it may be evolutionarily advantageous and achievable to recruit different enzyme families to perform ene-reduction.

SpiR and SpiE strongly cooccur across genomes and are encoded in a single operon in some species, indicating that they function as a biochemical pair. SpiR, which isomerizes and oxidizes cholesterol, is necessary to form the conjugated cholestenone intermediate for SpiE, acting as a redox pair. Given the abundance of cholesterol entering the colon and its potential as an electron acceptor for anaerobic respiration, the strong coevolution and narrow phylogenetic distribution of *spiR* and *spiE* are consistent with selection on this operon as a niche-defining metabolic capability that confers a competitive advantage in the cholesterol-rich gut environment.

The identification of SpiE and SpiR completes the microbial *spi* enzymatic pathway from cholesterol to coprostanol, enabling future mechanistic studies linking microbial cholesterol metabolism to host cholesterol levels. Consistent with this pathway’s relevance across human populations, SpiE abundance was strongly correlated with fecal coprostanol levels across gut microbiome cohorts and within individuals on antibiotics, suggesting SpiE is a robust biomarker of coprostanol-producing capacity in the human gut. These enzymes possibly allow gut microbes to influence host cholesterol levels, and probiotic options of cholesterol-metabolizing bacteria have been already explored in various contexts^36,37^. However, more in-depth *in vivo* studies and serum cholesterol measurements are required to assess whether there is a causal effect of cholesterol-metabolizing gut bacteria and host serum cholesterol levels. Serum cholesterol is also influenced by a wide variety of factors, including diet and host genetics. The identification of the *spi* pathway lays the groundwork for future studies to establish a potential causal effect of gut microbial cholesterol metabolism on serum cholesterol levels.

## Methods

### Identification of a candidate cholestenone 5β ene-reductase

All representative genomes of the family Acutalibacteraceae were downloaded from the Genome Taxonomy Database (GTDB) and annotated using Prokka (version 1.14.6)^38,39^. All sequences were annotated using InterProScan (version 5.72-103.0)^40^. OrthoFinder v3.1.0 was used to group the predicted protein sequences from each genome into orthogroups with an e-value cutoff of 1e-100^41^. We used our previous list of SpiR to define presence and absence in Acutalibacteraceae genomes (PMID: 41980965, Supplementary Data 2)^6^, and used Evolink to identify orthogroups associated with SpiR with default parameters^22^. Briefly, Evolink calculates an index that measures phenotype-genotype associations using Faith’s phylogenetic diversity of species with or without the genotype based on the presence of the phenotype, and then computes a significance threshold based on Isolation Forest detection of Evolink scores deviating from the background distribution^22^. We detected 4 out of 66,866 orthogroups significantly positively associated with SpiR. Only one out of four was annotated with putative oxidoreductase function (IPR051783, NAD(P)-dependent oxidoreductase-like) by InterProScan. This candidate was further investigated as putative cholestenone 5β ene-reductase.

### Profiling the taxonomic distribution of SpiE

Using the previously annotated GTDB protein sequences, we constructed an initial profile for SpiE using manually curated 492 SpiE homologs within Acutalibacteraceae, excluding abnormally short (<295 amino acids; 25 total) or long (>600 amino acids; one) sequences. These sequences were used to build a multiple sequence alignment (MSA) with MUSCLE (version 5.1)^42^. The resulting alignment was used to build a profile HMM with HMMER (version 3.4)^43^. The HMM was used to identify SpiE homologs in all GTDB sequences annotated with SSF51735 using hmmsearch. We set a bitscore cutoff of 525. We then removed a sequence for being less than 295 amino acids and two for being greater than 500 amino acids, resulting in 485 sequences.

The species tree of Acutalibacteraceae was obtained as a subset of the NCBI-GTDB bacteria tree using gotree (version 0.5.1) and the tree was collapsed to the genus level using ETE3 (version 3.1.2)^44,45^. Genera with fewer than 5 representative genomes were pruned. The resulting tree, including SpiE and SpiR prevalence per genus, was visualized using iTOL (version 7.6)^46^.

### Phylogenetic analysis of SDR ene-reductases

All SwissProt (accessed May 12, 2026) sequences annotated as SSF51735 superfamily enzymes classified as oxidoreductases (EC 1.-.-.-) were downloaded^47^, and sequences with an annotation score < 4 were removed. All sequences with the same gene name were collapsed, with the representative being the sequence with the highest annotation score or length closest to the median. mmseqs2 (version 2-13-45111-219-gaabc78c) was used to remove redundant sequences (--min-seq-id 0.7 -c 0.9)^48^. The resulting sequences were aligned using MUSCLE (version 5.1)^42^. Columns with >95% gaps were removed with Goalign (version 0.3.7)^44^. A phylogeny was inferred from the resulting trimmed alignment using raxml-ng (version 2.0.1), with 1,000 bootstraps^49^.

### Structural prediction, analysis, and catalytic residue prediction

The protein structure for SpiE was predicted using AlphaFold3 using default parameters^50^. During prediction, NADH (CID: 439153), cholestenone (CID: 91477) were specified as ligands. AlphaFold3 co-folds protein and ligands, so the ligand conformations were predicted jointly with the structure. The SpiE and *Digitalis lanata* progesterone 5β-reductase (PDB id: 2V6G) structures were aligned^25^, and the TM-scores were calculated using TM-align (v20220412)^51^. Structural homologs in the Protein Data Bank (PDB) were identified using the Foldseek web server (accessed August 14, 2026)^52^.

We calculated residue conservation with the goalign compute pssm command (v0.4.1)^44^, using the protein alignment of 485 homologs of SpiE. This command computes a Position-Specific Scoring Matrix (PSSM), which contains the probability of observing a specific residue at each position in the alignment. We inspected the residues conserved across all 485 homologs in the predicted structure, selecting those within 5 Å of either the carbonyl group or the alkene of cholestenone as putative catalytic residues. Protein sequence conservation was visualized using logomaker v0.8.7^53^.

### Profiling the relative abundance of SpiE in stool samples

We downloaded samples from leukemia (AML, n=13), liver disease (LD, n=153)^54,55^, ICU (MICU, n=52)^56^, and liver transplant (LT, n=56)^57^ cohorts, which had paired metagenomics and metabolomics data. We processed the metagenomic reads by trimming the adapter sequences using TrimGalore (v2.2.0) with default settings^58^, then mapping to a human reference (T2T-CHM13v2.0) to identify potential contaminants. Human reference-aligned reads were removed via Samtools (v1.23)^59^. We then aligned the reads to the SpiE reference database, built with the 485 SpiE homologs mentioned previously, using bowtie2 (v2.5.3)^60^. The number of reads mapped to SpiE per sample was normalized by dividing by the total reads and multiplying by one million to obtain counts per million (CPM). We removed samples with less than one million total reads after processing. SpiE homologs were considered present in a sample if the CPM was greater than or equal to 1 and metabolites present if the normalized area under the curve was greater than 0.1. Spearman’s correlation coefficients for SpiE CPM and coprostanol AUC were calculated using the cor.test function in R (v4.5.0).

### Taxonomic profiling of SpiE-encoding taxon in antibiotics-treated patients

We taxonomically profiled the aforementioned paired-end metagenomic reads using Kraken2 (v2.1.2) using GTDB release 226 reference genomes, generated using the Struo2 pipeline, which automates downloading GTDB genomes and Kraken2 database creation^61,62^. Reads were classified with a confidence threshold of 0.1. To estimate microbial abundances across multiple taxonomic ranks, we used Bracken (v2.8)^63^, which adjusts Kraken2 classifications by redistributing reads based on k-mer frequency models. Abundances were normalized by sequencing depth and reported as fractions of total classified reads. Final outputs included taxonomic identifiers and relative abundances at all hierarchical levels, from domain to species. SpiE-encoding taxa were defined as taxa encoding the aforementioned 485 SpiE homologs.

### Cloning and transformation

The gene *spiE* was synthesized using Genscript (GeneScript, Piscataway, NJ, USA) and the plasmid backbones used was pET28a(+) (Addgene, 69864-3). Construct-specific cloning strategies were selected based on the expression needs and compatibility with downstream applications. The constructs were transformed into chemically competent *E. coli* strains according to the manufacturer’s instructions: NEB T7 Express lysY/Iq (NEB, C3013I) for the pET28a(+) vector. Transformed cells were plated on LB agar containing 50 μg/mL kanamycin. Colonies with the recombinant plasmids were verified using Oxford Nanopore sequencing (Plasmidsaurus).

### Mutant generation

The *Eubacterium coprostanoligenes* ATCC 51222 *spiE* gene in T7 Express *lysY E. coli* was used as the template for site-directed mutagenesis. Primers were designed in SnapGene for D77A and Y138A, with annealing regions flanking the mutation (**Supplementary Data 4**). Mutants were amplified from plasmid templates, verified by gel electrophoresis, digested with DpnI (NEB, R0176S) to remove parental DNA, and transformed into T7 Express *lysY E. coli*. R79A and S118A mutants were generated using the Q5 Mutagenesis Kit (NEB, E0554S) according to the manufacturer’s protocol. Recombinant plasmids were verified by Oxford Nanopore sequencing (Plasmidsaurus).

### Whole cell assays

Transformed *E. coli* strains were inoculated from glycerol stocks into 250 mL LB containing 50 µg/mL kanamycin and grown overnight at 32 °C. Cells were pelleted at 3,260 × *g* for 10 minutes and transferred to an anoxic chamber. Pellets were resuspended in 100 mL anoxic LB containing 10 µg/mL progesterone or pregnenolone, 50 µg/mL kanamycin, and 400 µM IPTG (GoldBio, 22110512481), then incubated anoxically at 37 °C for 48 hours. Cells were pelleted at 3,260 × *g* for 10 minutes, and steroids were extracted from the supernatant with three mL chloroform (Fisher Chemical, C298) using a separatory funnel. The chloroform phase was air-dried and the extract was resuspended in one mL methanol (Fisher Chemical, 232699).

### UHPLC-MS analysis of steroid hormones

The whole-cell assay samples resuspended in methanol were analyzed using a Bruker Maxis-II QTOF ultra-high-resolution Q-TOF mass spectrometer coupled with a Waters Acquity I-Class PLUS LC system. Liquid chromatography separation was performed on a Phenomenex Kinetex F5 100 Å LC Column (100 × 3 mm, 2.6 μm particle size) with mobile phase A (water with 0.1% formic acid) and mobile phase B (acetonitrile with 0.1% formic acid). The gradient program began with 10% B for 2 min, increased to 55% B over the next minute, and further increased to 65% B for 11 min. B was increased to 90% over the next minute, maintained at 90% B for 2 min, decreased to 10% over the course of 1 min, and maintained for 3 min. The column was maintained at 45 °C. The injection volume was 5 µL. Mass spectra were acquired under positive electrospray ionization with an ion spray voltage of 4500 V. The source temperature (dry gas) was set to 220 °C with a flow rate of 5.0 L/min. Mass spectrometric analysis was performed on a Bruker maXis II Q-TOF instrument controlled by Bruker otofControl (v5.2) and Bruker HyStar (v5.1.8.1). Extracted ion chromatograms were generated at theoretical [M + H]⁺ *m*/*z* values using ±0.05 Da tolerance for chromatographic profiling and 20 ppm for targeted detection. Data processing was performed in Python (v3.10.12) using pyOpenMS (v3.1.0), pyteomics (v4.7.5).

### Lysate Assay

Ten milliliters of overnight *E. coli* culture were inoculated into 500 mL LB containing 50 µg/mL kanamycin and grown at 32 °C to an OD600 of 0.5–0.6. Protein expression was induced with 400 µM IPTG, followed by overnight shaking at 25 °C. Cells were pelleted at 3,260 × *g* for 10 min and stored at −20 °C. Pellets were thawed on ice for 30 min and resuspended in pH-adjusted lysis buffer (100 mM NaPO4, 350 mM NaCl, 10 mM BME, pH 6.5) containing lysozyme (Research Products International, L38100), 100 µL BME (G Biosciences, BC98), and protease inhibitor (Sigma-Aldrich, 04693124001). Cells were sonicated at 4 °C using a Branson 550 sonicator at 40% amplitude (15 s on/30 s off) for 2 min total. Lysates were clarified at 14,000 × *g* for 10 min.

The supernatant was incubated anaerobically with 100 µL of 100 mM cholesterol (IBI Scientific, IB15721), 10 µL of 100 mM NAD^+^ (Sigma-Aldrich, 10127981001) in the same lysis buffer, and 10 µL of 100 mM NADP^+^ (Sigma-Aldrich, 10128031001) in buffer at 4 °C for 24 hours and transferred to 37°C for 2 hours. Prior to addition to the lysate, cholesterol and cholestenone were solubilized using methyl-β-cyclodextrin (Sigma Aldrich, C4555). The products were extracted using chloroform and resuspended in methanol for GC-MS analysis. This same process was repeated with 100 µL of 100 mM cholestenone (Ambeed Inc., 601-57-0), 10 µL of 100 mM NADH (Cayman Chemical, 1949720-50-6), and 10 µL of 100 mM NADPH (Cayman Chemical, 2646-71-1).

### Targeted GC–MS analysis of cholesterol metabolites from heterologous expression assays

#### Sample preparation

Samples were kept on ice throughout processing. A 200-µL aliquot of the methanolic extract was transferred to a GC vial and dried under a gentle stream of nitrogen.

Dried extracts were derivatized by resuspension in 80 µL of freshly prepared methoxyamine hydrochloride in pyridine (20 mg/mL), mixed vigorously for 5 min and incubated for 2 h at 50 °C (methoximation), followed by addition of 80 µL of MSTFA and incubation for a further 1 h at 50 °C (trimethylsilylation) immediately prior to GC–MS analysis. Ribitol served as an extraction/derivatization control and was not used for quantification.

#### Calibration standards

Individual analytical standards were prepared as 10 mg/mL stocks in chloroform:methanol (2:1, v/v) and combined into a working mixture at 100 µg/mL, which was serially diluted (alternating 1:1 and 1:5) to yield an eight-point external calibration series (0, 0.1, 0.5, 1, 5, 10, 50 and 100 µg/mL). A 200-µL aliquot of each level was dried and derivatized identically to the samples. Each acquisition batch included its own full calibration series, interspersed reagent blanks, and pooled-QC injections.

#### GC–MS analysis

Derivatized extracts (1 µL) were injected in split mode (10:1 split ratio; split flow 15 mL/min; septum purge 1 mL/min) with the inlet at 250 °C on an Agilent 8890 GC coupled to an Agilent 7200B quadrupole time-of-flight (Q-TOF) mass spectrometer (MassHunter GC/MS acquisition). Separation was performed on an Agilent VF-5ms capillary column (30 m × 0.25 mm i.d. × 0.25 µm film; CP8944) with helium carrier gas at a constant flow of 1.5 mL/min. The oven was held at 185 °C for 1 min, ramped at 10 °C/min to 310 °C (held 3 min), then at 25 °C/min to 340 °C (held 7 min), for a total run time of 24.7 min; the transfer line was held at 280 °C. The mass spectrometer was operated in electron ionization (EI) mode at 70 eV with the ion source at 230 °C; full-scan (MS1) spectra were acquired in centroid mode over m/z 50–500 at 5 spectra/s following a 2-min solvent delay.

#### Data processing and quantification

Raw instrument files (.D) were converted to mzML (msconvert/ProteoWizard), and each analyte was extracted as a full-scan chromatogram at its retention time within a per-compound tolerance window, empirically re-centered for each acquisition batch to account for instrument drift, using two characteristic ions per compound at a mass tolerance of approximately 0.05–0.07 Da. Quantification used log-transformed external calibration curves (linear or quadratic fit selected per compound) built from the eight-point calibration series, with limits of detection and quantification derived directly from each curve’s fitted slope and residual noise at the lowest standard. Peak extraction, calibration fitting, and concentration calculation were carried out with an in-house Python/R pipeline (Arome Science Inc., https://www.arome-science.com/metabolomics/scaling-gc-ms-data-analysis/) run in targeted, full-scan (MS1) mode. Each batch was quantified against its own calibration series; reported concentrations are µg/mL of extract and are not normalized to sample mass or volume.

### Clinical Sample Collection

Patients from four clinical studies were prospectively recruited for stool sample collection. Patients were included based on disease state present at the time of hospitalization (AML, LD, LT) or location of admission (MICU). Informed consent was obtained for all patients in accordance with the Helsinki convention, and under the supervision of the University of Chicago Institutional Review Board, (IRB 20-0163, IRB 20-1384, IRB 20-1102, IRB 21-0327, IRB 23-1365). All samples were stored at 4℃ for <24 hours before being aliquoted and stored at -80℃ prior to metabolomic analysis.

### Coprostanol measurement from human fecal samples

Coprostanol from human fecal samples in the AML, LD, LT and MICU cohorts were measured by the Duchossois Family Institute Microbiome Metabolomics facility at the University of Chicago. Metabolites were extracted with the addition of organic solvent (80% methanol spiked with heavy labeled internal standards) to pre-weighed fecal samples at a ratio of 100 mg of material per mL of extraction solvent in beadruptor tubes (Fisherbrand; 15-340-154). Samples were homogenized at 4° C on a Bead Mill 24 Homogenizer (Fisher; 15-340-163), set at 1.6 m/s with 6 thirty-second cycles, 5 seconds off per cycle. Samples were then centrifuged at -10°C, 20,000 x *g* for 15 min and the supernatant was used for subsequent metabolomic analysis. All metabolites are compared to authentically purchased standards for retention time, *m/z* and fragmentation pattern on the same instrument. Metabolites were analyzed using GCMS with electron impact ionization. The metabolite extract (100 µL) was added to a mass spectrometry autosampler vial (Microliter; 09-1200) and dried down completely under nitrogen stream at 30 L/min (top) 1 L/min (bottom) at 30 °C (Biotage SPE Dry 96 Dual; 3579M). To dried samples, 50 µL of freshly prepared 20 mg/mL methoxyamine (Sigma; 226904) in pyridine (Sigma; 270970) was added and incubated in a thermomixer C (Eppendorf) for 90 min at 30°C and 1400 rpm. After samples are cooled to room temperature, 80 µL of derivatizing reagent (BSTFA + 1% TMCS; Sigma; B-023) and 70 µL of ethyl acetate (Sigma; 439169) were added and samples were incubated in a thermomixer at 70 °C for 1 hour and 1400 rpm. Samples were cooled to RT and 400 µL of Ethyl Acetate was added to dilute samples. Turbid samples were transferred to microcentrifuge tubes and centrifuged at 4 °C, 20,000 x *g* for 15 min. Supernatants were then added to mass spec vials for GCMS analysis. Samples (1 µL injection) were separated and analyzed using a HP-5MSUI column (30 m x 0.25 mm, 0.25 µm; Agilent Technologies 19091S-433UI) coupled to a GC-EI-MS (Agilent 7890A GC system, Agilent 5975C MS detector) with the following oven ramp parameters: 1 min hold at 60 °C, 16 °C per min up to 300 °C with a 7 min hold at 300 °C. Inlet temperature was 280 °C and transfer line was 300 °C. Data analysis on raw, proprietary files (.d) was performed using MassHunter Quantitative Analysis software (version B.10, Agilent Technologies). Normalized peak areas were calculated by dividing raw peak areas of targeted analytes by averaged raw peak areas of internal standards.

### Protein purification

SpiE-expressing *E. coli* strains were grown overnight at 37 °C in LB containing 50 µg/mL kanamycin. A 7.5 mL overnight culture was used to inoculate 750 mL LB and grown to an OD_600_ of ∼0.4. Expression was induced with 400 µM IPTG, followed by overnight incubation at 16 °C. Cells were harvested at 3,260 × *g* for 10 minutes, stored at −20 °C for two hours, thawed on ice, and resuspended in lysis buffer (0.1 M NaPO_4_, 0.35 M NaCl, 5 mM BME, 5% w/v glycerol, pH 6.5) supplemented with lysozyme, benzonase (Sigma-Aldrich, E8263-5KU), and protease inhibitor.

Cells were sonicated at 4 °C using a Branson 550 sonicator at 40% amplitude (15 s on/30 s off) for two minutes total. Lysates were clarified at 14,000 × *g* for 10 minutes, filtered through a 0.22 µm PES filter (Celltreat, 229746), and buffer-exchanged into lysis buffer. SpiE was purified using a HisTrap Excel column (Cytiva, 17371205) and eluted with a linear gradient of lysis buffer containing 0.2 M imidazole. The protein was concentrated again into the lysis buffer using 10 kDa MWCO Amicon Ultra filters (Millipore Sigma, MPE010025). SpiE expression and purification were confirmed by SDS-PAGE.

### Binding characterization

SpiE was concentrated to 100 µM in 0.1M NaPO4, 0.35M NaCl, 5mM BME, 5% w/v glycerol at a pH:6.5 supplemented with 5% DMSO (AG Scientific, C-1019) v/v. The ligands tested were resuspended and prepared in an identical buffer. Cholestenone (TargetMol, 601-57-0) required additional preparation to ensure solubility, it was suspended in 100% DMSO at 10 mM concentration and shaken at RT for one week before being diluted to their working concentration of 10 µM.

ITC experiments were performed at 25 °C using a TA Instruments Nano ITC instrument. Cholestenone and progesterone (Millipore Sigma, 57-83-0) were assayed using inverse titrations with SpiE in the syringe and the ligand in the cell. SpiE was injected into 3.1 µL aliquots at 120-second intervals with a stirring speed of 150 RPM. Heats of dilution were subtracted, and the data were analyzed using NanoAnalyze software (TA Instruments, v3.7.0). Binding curves were generated to calculate the enthalpy change (ΔH), entropy change (ΔS), stoichiometry (n), and dissociation constant (Kd) based on the independent binding model.

## Author Contributions

X.J., B.H., A.K.J., and G.A. contributed to the conceptualization of the study. X.J., B.H., A.K.J., and G.A. developed the methodology and performed validation. Formal analysis was conducted by A.K.J., G.A., M.K., M.R.G., T.B., and X.J., while investigation was carried out by A.K.J, G.A., M.K., M.R.G., T.B., A.M.C., A.M.S., M.M., C.L., M.O., K.W., B.P., O.O., S.H.L, Y.L., B.H., and X.J., and data curation was performed by A.K.J., M.K., X.J., and G.A. The original draft of the manuscript was written by A.K.J., G.A., and M.K., with subsequent review and editing by S.H.L, G.A., M.K., M.R.G., A.K.J, A.M.S., X.J., and B.H. Visualization efforts were led by X.J., A.K.J., G.A., M.K., and T.B. Supervision was provided by S.H.L, Y.L., B.H. and X.J.

## Acknowledgements

This study utilized the computational resources of the NIH HPC Biowulf cluster (http://hpc.nih.gov). GC-MS analysis and data processing were performed by Arome Science Inc. and the Duchossois Family Institute.

A.K.J, M.K., G.A., and X.J. are supported by the Intramural Research Program of the NIH, National Library of Medicine. B.H., and M.R.G. are supported by R35-GM155208. S.H.L is supported by grants R21-AI199298 and R35-GM146969. This research was supported in part by the Intramural Research Program of the National Institutes of Health (NIH). The contributions of the NIH authors are considered works of the United States Government. The findings and conclusions presented in this paper are those of the authors and do not necessarily reflect the views of the NIH or the U.S. Department of Health and Human Services.

## Data and materials availability

The authors confirm that the data supporting the findings of this study are available within the article and its supplementary materials. All genomic data is available at the GTDB. All reads mapped to our SpiE database are publicly available and the accession numbers can be found in **Table S3**.

