## Supporting Information for "SpiE is a distinct gut microbial 4-cholesten-3-one 5β-reductase that acts with SpiR to convert cholesterol to coprostanol"

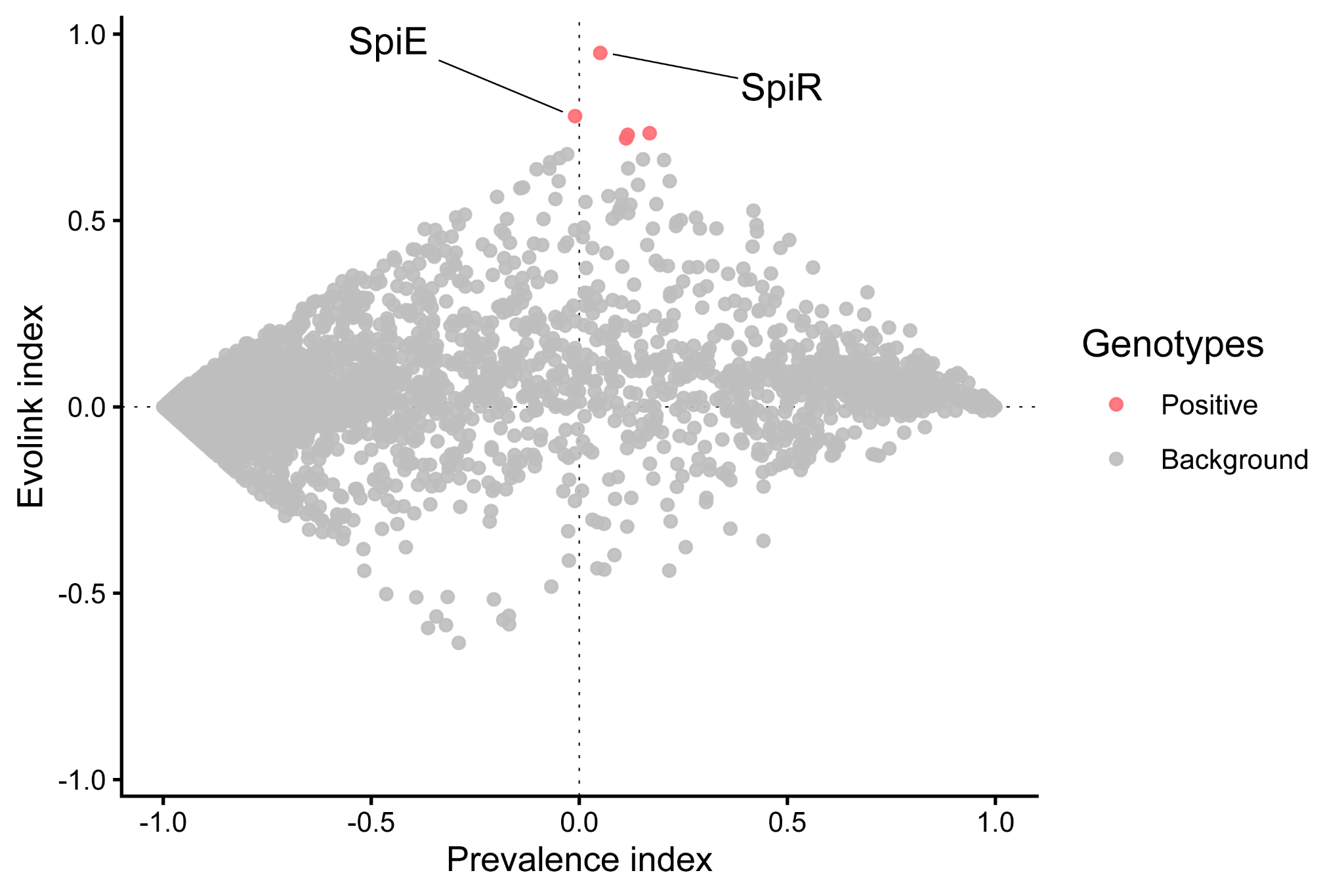

**Figure S1:** Scatterplot of the Evolink index (value from -1.0 to 1.0, where 1.0 represents perfect correlation to SpiR and -1.0 indicating the opposite) and the prevalence index for all Acutalibacteraceae gene families. Each point represents a single gene family. Gene families significantly correlated to SpiR are colored red.

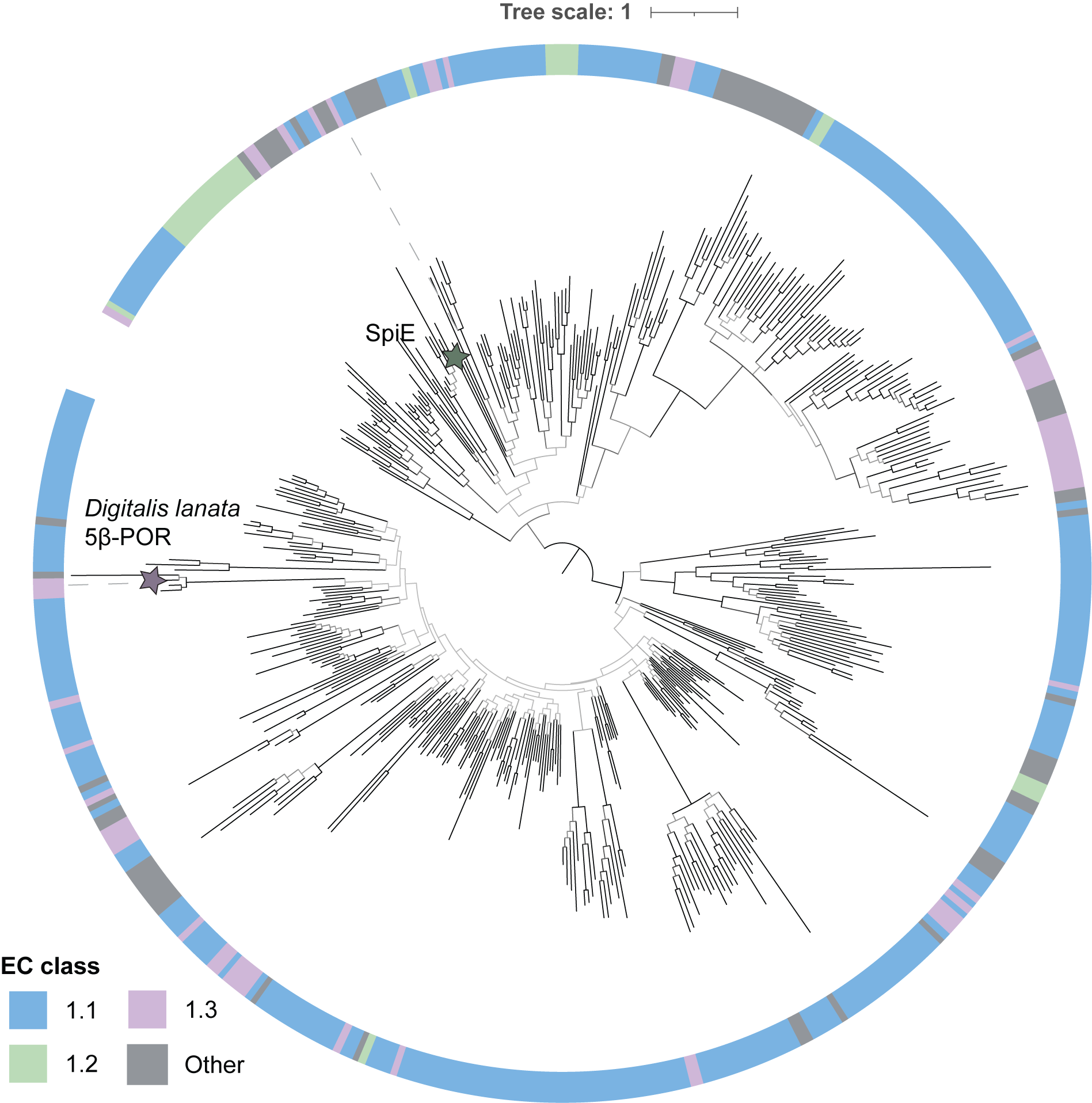

**Figure S2:** Maximum likelihood protein phylogeny of curated SwissProt SDR oxidoreductases showing that SpiE belongs to a distinct enzyme family. The branches are colored by bootstrap support values, with darker branches indicating higher values. SpiE and *Digitalis lanata* progesterone 5β-reductase (5β-POR) are labeled on the tree. The outer ring is color-coded by EC number (1.1 acting on CH-OH group, 1.2 acting on the aldehyde or oxo group, 1.3 acting on CH-CH group, and other 1.-.-.-).

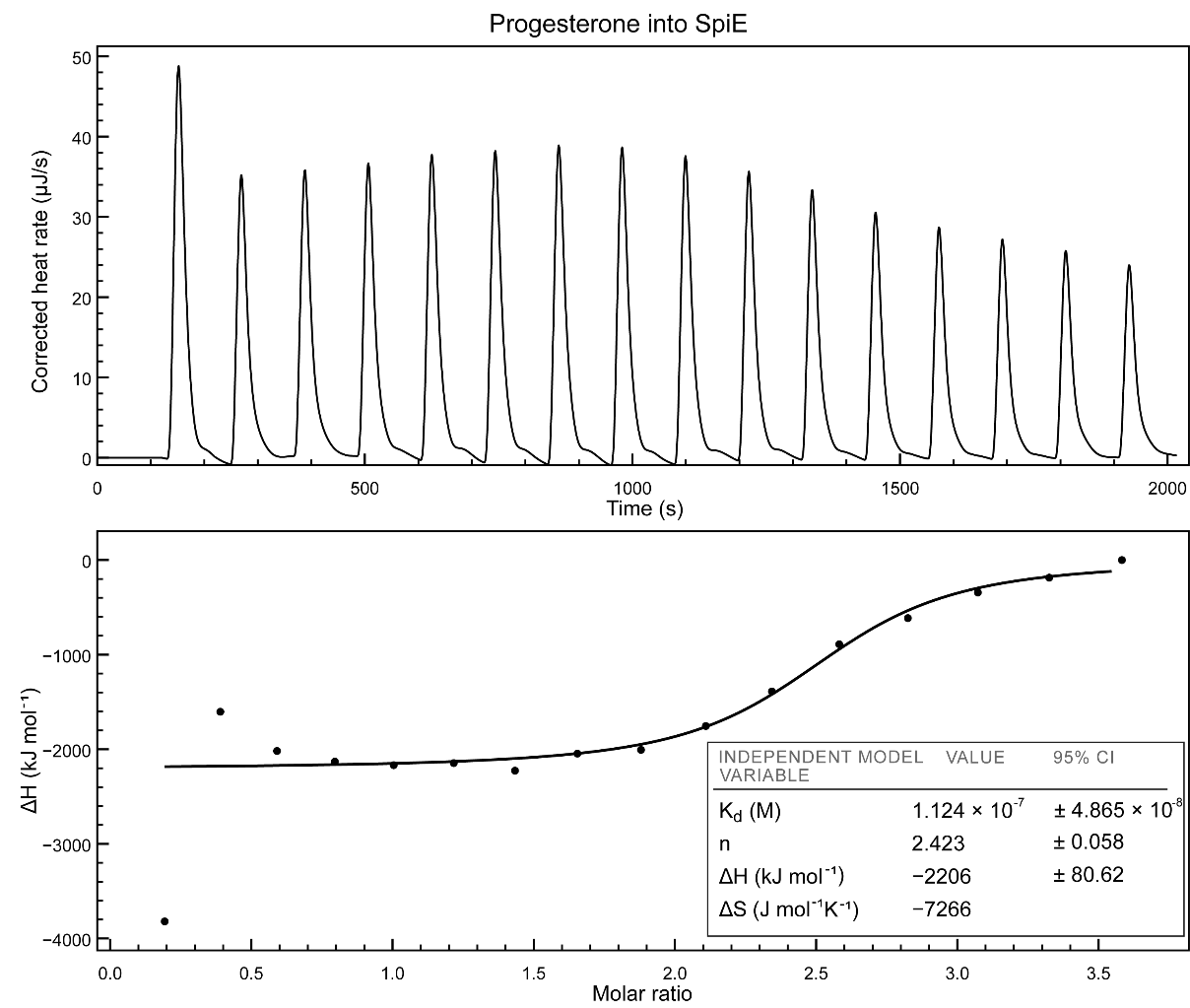

**Figure S3: SpiE binding to progesterone:** Binding of SpiE to progesterone as measured by ITC. Raw data is shown in the upper graph of each panel, integrated binding isotherms with their independent binding model fitting are shown in the lower panel. Recombinant SpiE was placed in the calorimeter cell, and the experiment was performed at 25 °C with successive 3.1 µL injections of progesterone. The binding of the trace was performed using a 100 µM progesterone and 10 µM SpiE.

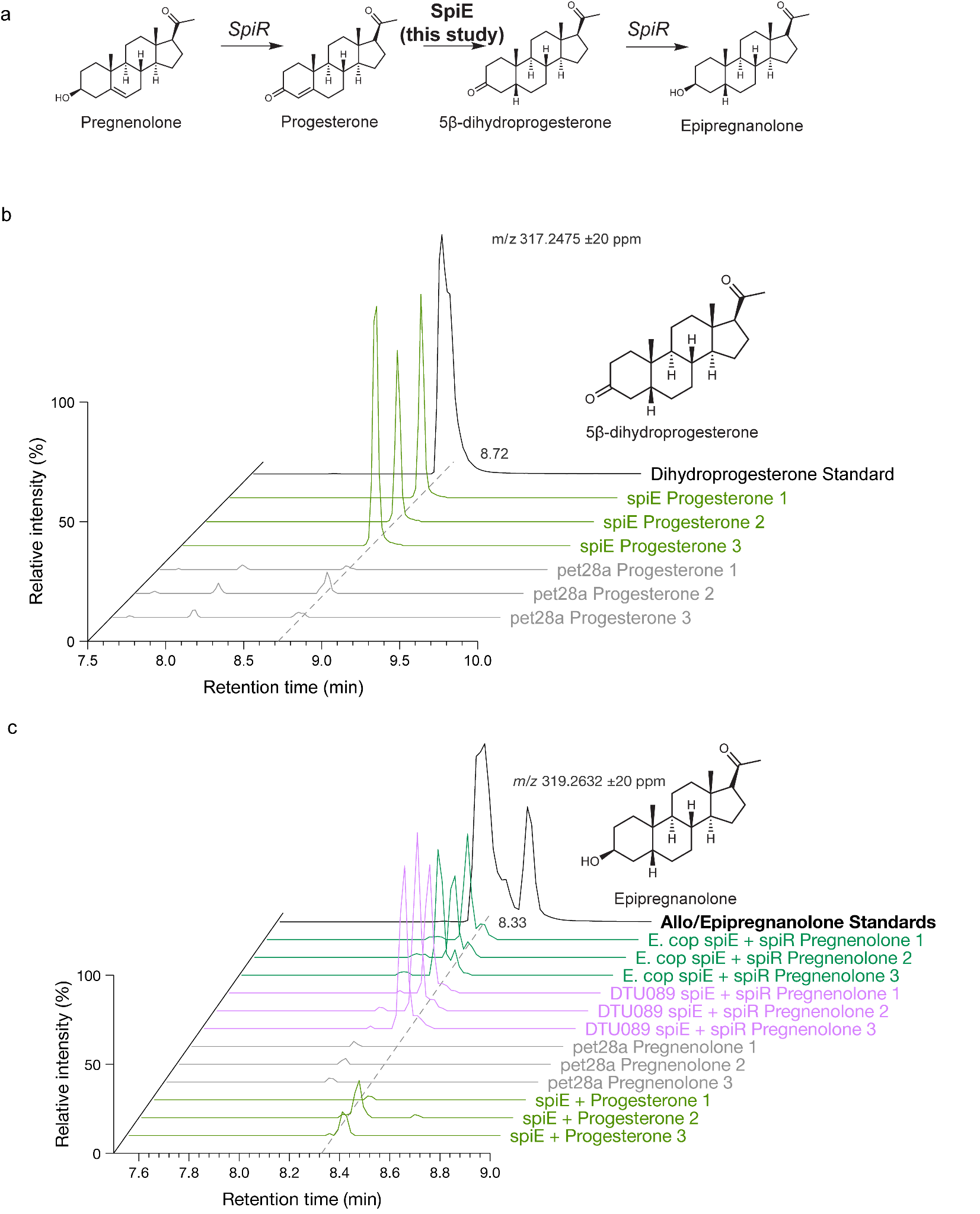

**Figure S4: SpiE activity on progesterone.** a) Reaction scheme of SpiE and SpiR on pregnenolone. SpiE performs the reduction of progesterone into dihydroprogesterone. b) Chromatogram of heterologous expression of SpiE with progesterone, showing production of dihydroprogesterone. c) Chromatogram of heterologous expression of SpiE, or SpiE and SpiR (from *E. coprostanoligenes* and *DTU089 sp.*), showing that SpiR and SpiE together produce epipregnanolone, while SpiE alone does not.

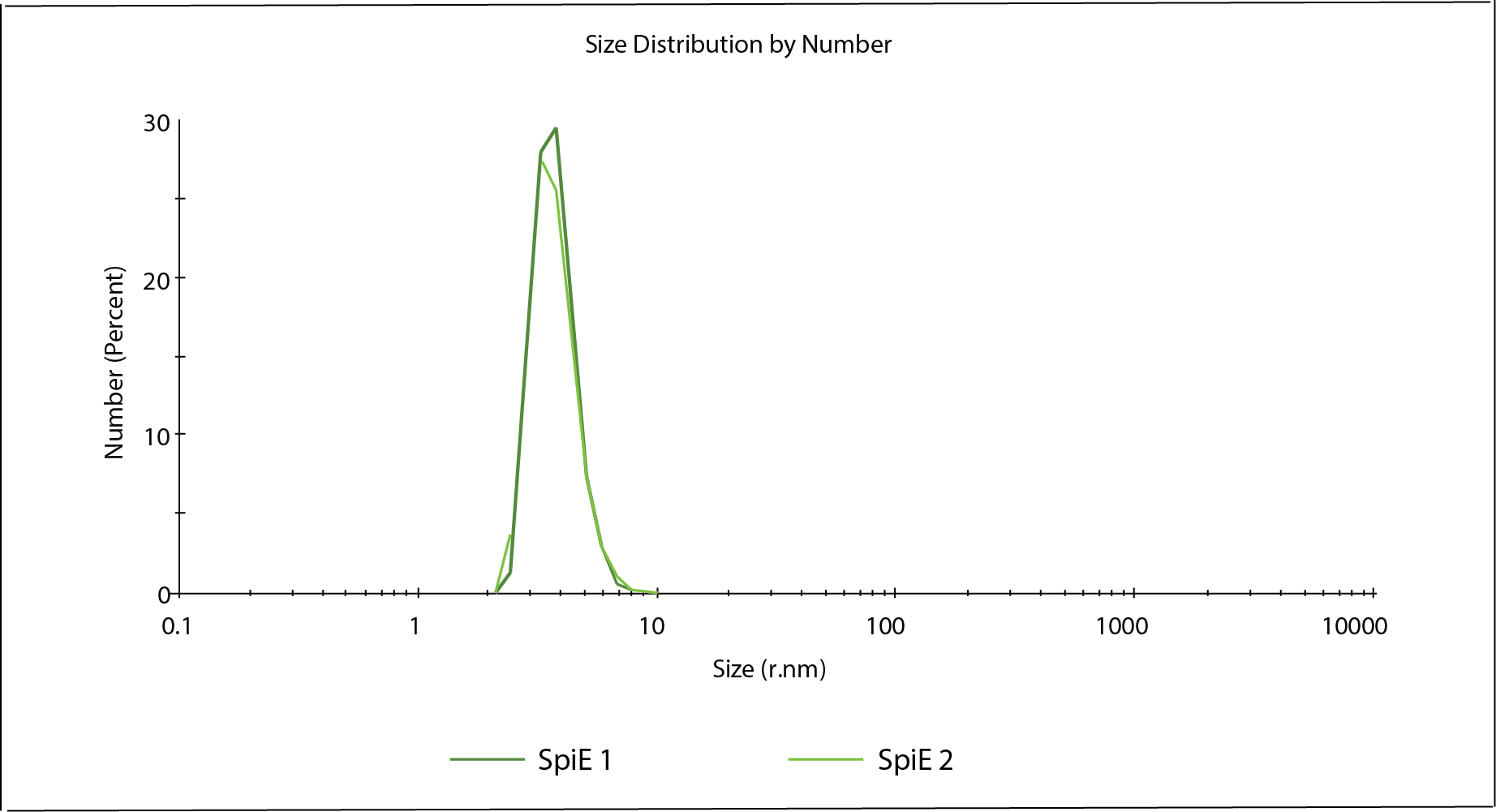

**Figure S5. SpiE is a homodimer in solution:** The hydrodynamic radius of the enzyme is 3.2 nm reported in nanometers on the x-axis. The y-axis indicates the relative abundance of each size population. The size distribution of the sample is shown by number to correct for the overrepresentation of large aggregates that scatter light more intensely even though they are present at reduced amounts. SpiE 1(dark green) and SpiE 2 (light green) are the duplicate technical measurements of the enzyme. Each final measurement displayed, is the average of 15 separate measurements taken over the span of 120 seconds.

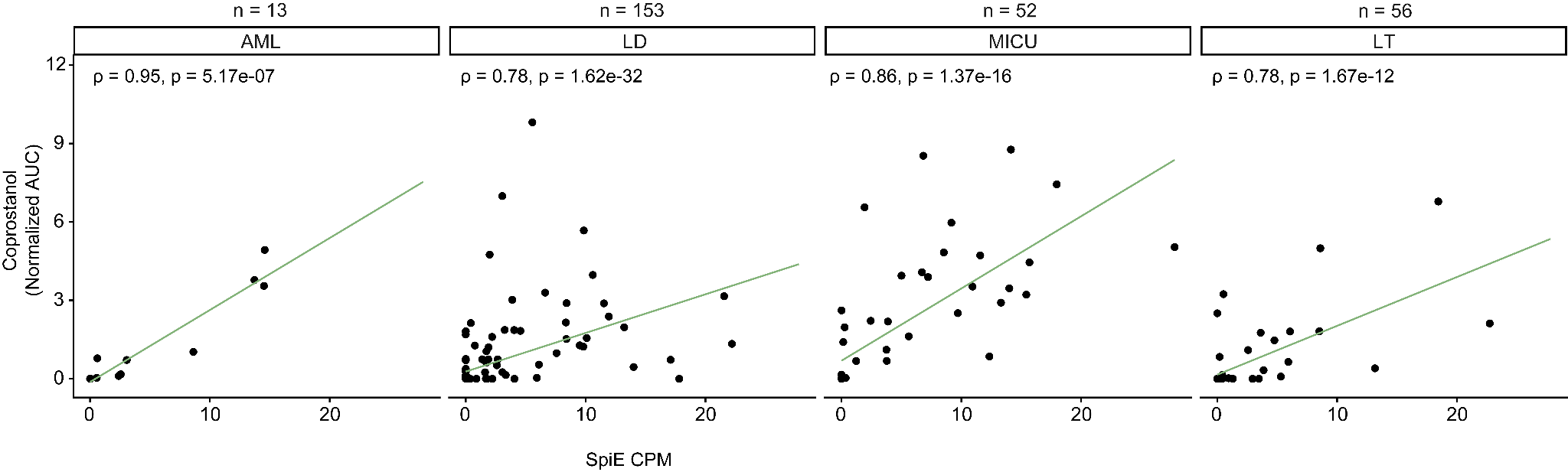

**Figure S6.** Association between SpiE homolog abundance in CPM and coprostanol abundance in normalized area under the curve (AUC) for paired human metagenomic and metabolomic cohorts. The data were fitted with a linear regression line. ρ represents the Spearman’s correlation coefficient.

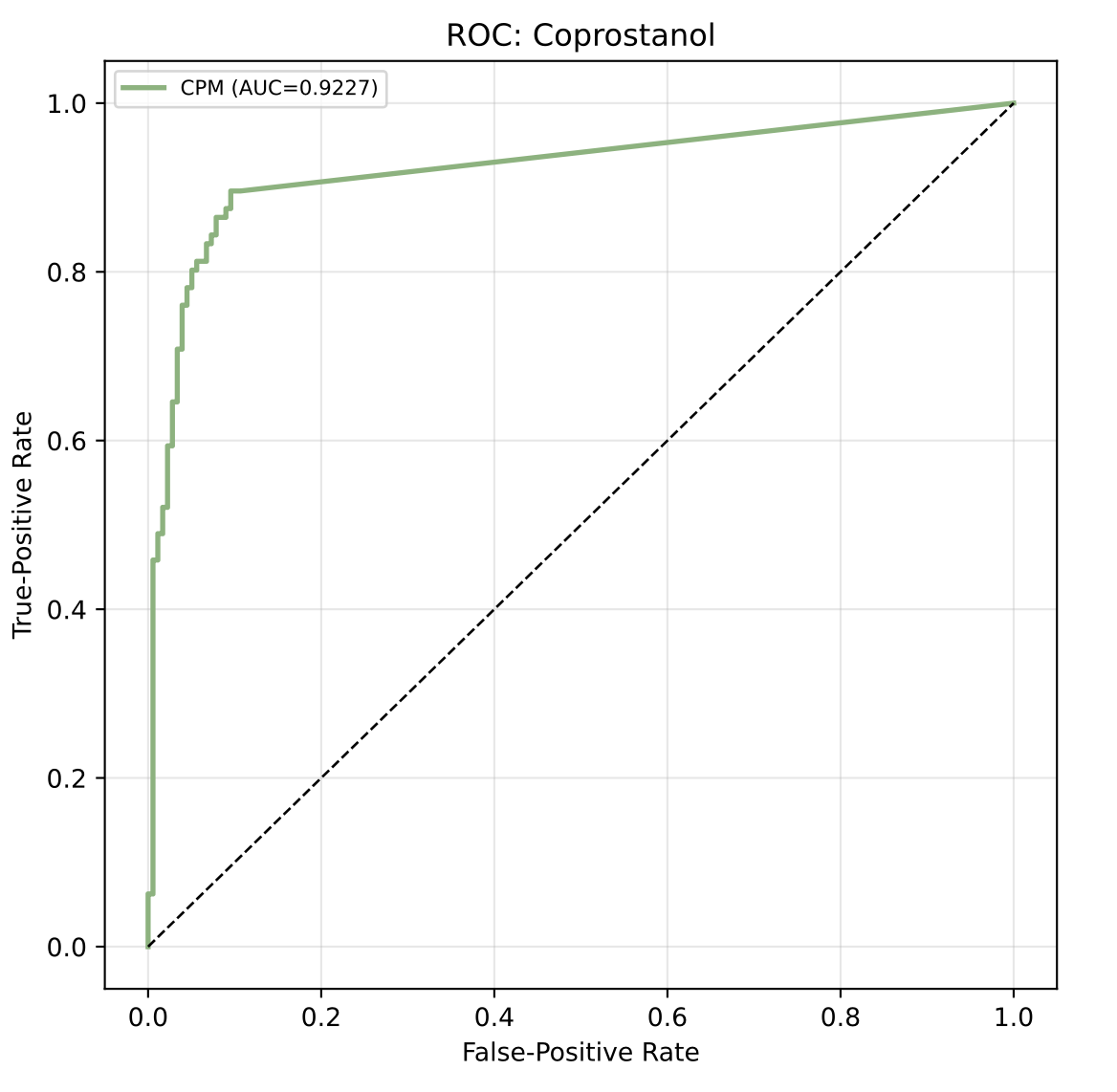

**Figure S7.** Receiver operating characteristic curve (ROC) for the prediction of coprostanol detection in stool samples from all four paired metagenomic and metabolomic cohorts (AML, LD, MICU, and LT) based on SpiE homolog abundance in CPM. The cutoff for coprostanol presence was >0.1 normalized area under the curve (AUC). The ROC curve and AUC values were calculated using the Python package scikit-learn (v1.9.0).
